# From Policy to Practice: Tracking an Open Science Funding Initiative

**DOI:** 10.1101/2023.02.27.530238

**Authors:** Kristen Ratan, Sonya B. Dumanis, Souad McIntosh, Hetal V. Shah, Matt Lewis, Timothy H. Vines, Randy Schekman, Ekemini A Riley

## Abstract

This is a critical moment in the open science landscape. Over the past few years there has been growing momentum to improve open research policies and require grantees to share all research outputs, from datasets to code to protocols, in FAIR (findable, accessible, interoperable and reusable [FAIR]) repositories with persistent identifiers attached. The Aligning Science Across Parkinson’s (ASAP) initiative has made substantial investments in improving open science compliance monitoring for its grantees, requiring grantees to update their manuscripts if not all research outputs have been linked in the initial manuscript version. Here, we evaluate ASAP’s effectiveness in improving research output sharing for all articles processed through the ASAP compliance workflow between March 1, 2022, and October 1, 2022. Our ultimate goal in sharing our findings is to assist other funders and institutions as they consider open science implementation. By normalizing the open science and compliance process across funding bodies, we hope to simplify and streamline researcher, institutional, and funder workflows, allowing researchers to focus on science by easily leveraging resources and building upon the work of others.

## Introduction: open research momentum

The quest for a COVID vaccine showed the world that science can advance at a rapid pace, efficiently and effectively, with researchers around the globe collaborating, iterating, and building upon each other’s work. This collaborative effort was an exception to the usual pace of research, often defined by siloed communities and inaccessibility to research outputs. The lack of data sharing, accurate citing of resources and protocols, and varying publication standards have created a fractured and incomplete research communication ecosystem. In the digital age, connecting the vast network of shared knowledge will only become more difficult as the sheer volume of research outputs and findings grows.

Over the past few years there has been growing momentum to address these concerns and improve open research policies. Open research involves the sharing of all research outputs, from datasets to code to protocols, in FAIR (findable, accessible, interoperable and reusable [FAIR]) repositories with persistent identifiers attached. The 2021 [UNESCO Recommendation for Open Science](https://unesdoc.unesco.org/ark:/48223/pf0000379949.locale=en) and the 2022 [Budapest Open Access Initiative 20th anniversary recommendations](https://www.budapestopenaccessinitiative.org/boai20/) are two of many recent reports calling for open knowledge and output sharing.

Building on this momentum, in August 2022, the [US White House Office of Science and Technology (OSTP) released a memo](https://www.whitehouse.gov/ostp/news-updates/2022/08/25/breakthroughs-for-alldelivering-equitable-access-to-americas-research/) with guidance that all federally funded research articles be 1) open access and 2) include sharing of underlying datasets in public repositories. The new OSTP memo is likely a bellwether for policy shifts towards open research in the US.

This is a critical moment in the open science landscape and it is imperative that funders, institutions, and governments establish common policies, best practices, and infrastructure (Staunton, et al., 2021). Collective action to align on open science policies and practices needs to happen now, in order to reduce the cost and friction of adoption, as well as increase collaboration, reuse, and reproducibility among researchers (Gabelica, Bojčić, Puljak (2022); Haak, Greene and Ratan, 2020; Zariffa, Haggstrom, & Rockhold, 2021).

The [Aligning Science Across Parkinson’s (ASAP) initiative](https://parkinsonsroadmap.org/) launched in 2019 with the mission of accelerating the pace of discovery and informing the path to a cure for Parkinson’s disease through collaboration, research-enabling resources, and data sharing (Schekman and Riley, 2019). Integral to the initiative was the establishment of a progressive [open science policy](https://parkinsonsroadmap.org/open-access-policy/). In a scientific landscape that encourages competition, ASAP’s founding conviction is that more intentional collaboration leads to faster and better outcomes (Allen, OConnell, and Kiermer, 2019; CODATA Coordinated Expert Group, Berkman et al, 2020). To cultivate this goal, ASAP established a research culture that incentivizes early sharing and collaboration rather than highly competitive practices that focus on high impact publishing. (Alberts et al., 2014; Anderson, Ronning, De Vries and Martinson, 2007; Bilder, Lin and Neylon, 2015; National Academies of Sciences, Engineering, and Medicine, 2021; Skinner and Lippincott, 2020; UNESCO, 2021). The ASAP [Collaborative Research Network (CRN)](https://parkinsonsroadmap.org/research-network/), is an international, multidisciplinary, and multi-institutional network of collaborating investigators working to address high-priority basic science questions related to Parkinson’s disease pathology in a highly collaborative environment. The CRN currently funds 35 teams that are composed of 163 lead investigators from around the world, including 80+ institutions, 14 countries, 51 female principal investigators, and 53 early-career investigators. Each funded team is required to hire a project manager to facilitate open and collaborative research practices. A detailed blueprint of the CRN, templates, reports, and other resources has been published in the [ASAP Blueprint for Collaborative Open Science](https://zenodo.org/record/6979998#.Y_9RLD1BxPY). ASAP’s grantees are already compliant with the OSTP memo and the emerging requirements of other funders in the EU and around the globe.

## Born Compliant: Building the open research tool chain and best practices

Below are the main points covered in the OSTP memo and a description of ASAP’s approach through the CRN.

- **Immediate open access with no embargo**: ASAP requires immediate open access in addition to a mandatory CC-BY preprint at the time of (or before) article submission.
- **Dataset Sharing**: ASAP’s policy requires that all underlying research outputs (protocols, code, datasets) be posted to a FAIR repository at the time of the mandatory preprint. ASAP does not mandate which FAIR repository (Wilkinson, et al. 2016) to use. However, ASAP has established relationships with existing repositories that offer community functionality, such as Protocols.io and [Zenodo](https://zenodo.org/communities/asaphub/?page=1&size=20), to help grantees when recommendations are needed. Delays or omissions at the stage of pre-printing are tracked and must be rectified by the time of article publication.
- **Research Reusability**: The guidance emphasizes the need for machine-readability formats and also licensing that allows for reuse. ASAP grantees publish in traditional preprint servers and journals with appropriate metadata. In addition, CC-BY licensing is required for preprints and journal articles.
- **Metadata and Persistent Identifiers**: All research outputs from ASAP-funded research must have DOIs or other appropriate identifiers, such as RRIDs for material resources as well as adequate metadata. These identifiers must be appropriately linked in the manuscript. In addition, all grantees are required to have an ORCID (Open Researcher and Contributor ID) to be part of the collaborative research network (CRN)).

Here, we examine the effectiveness of ASAP policies and the lessons learned for other funders and institutions as they consider implementation. Normalizing the open science and compliance process across funding bodies will greatly simplify and streamline researcher, institutional, and funder workflows and allow researchers to focus on science by easily leveraging resources and building upon the work of others.

## Tracking Compliance: Conducting research on open research

With the joint objective of guiding, supporting and assisting CRN teams in understanding ASAP’s open science requirements and understanding teams’ baseline compliance, ASAP partnered with an AI startup, [DataSeer](https://dataseer.ai/), which has developed software that uses natural language processing and machine learning processes to identify and assess the research outputs in a manuscript. The software simultaneously tallies the quantity, citations, and sharing status of newly generated and existing datasets, code, software, protocols, and lab materials. A report is automatically generated, and assessed by a DataSeer curator, summarizing action items required for the article to meet compliance with ASAP policies. Finally, ASAP staff review the report and send it to the authors to make amendments. The submission, curation, and adjustment process are iterative, with the ASAP team providing continual feedback until compliance is achieved. An example template of the report was deposited in Zenodo (https://doi.org/10.5281/zenodo.7504034). Articles for DataSeer review are submitted to ASAP via each team’s project manager or through discovery of articles via OA.Report (RRID:SCR 023288). See Figure 1 for the workflow schematic.

**Figure 1:**
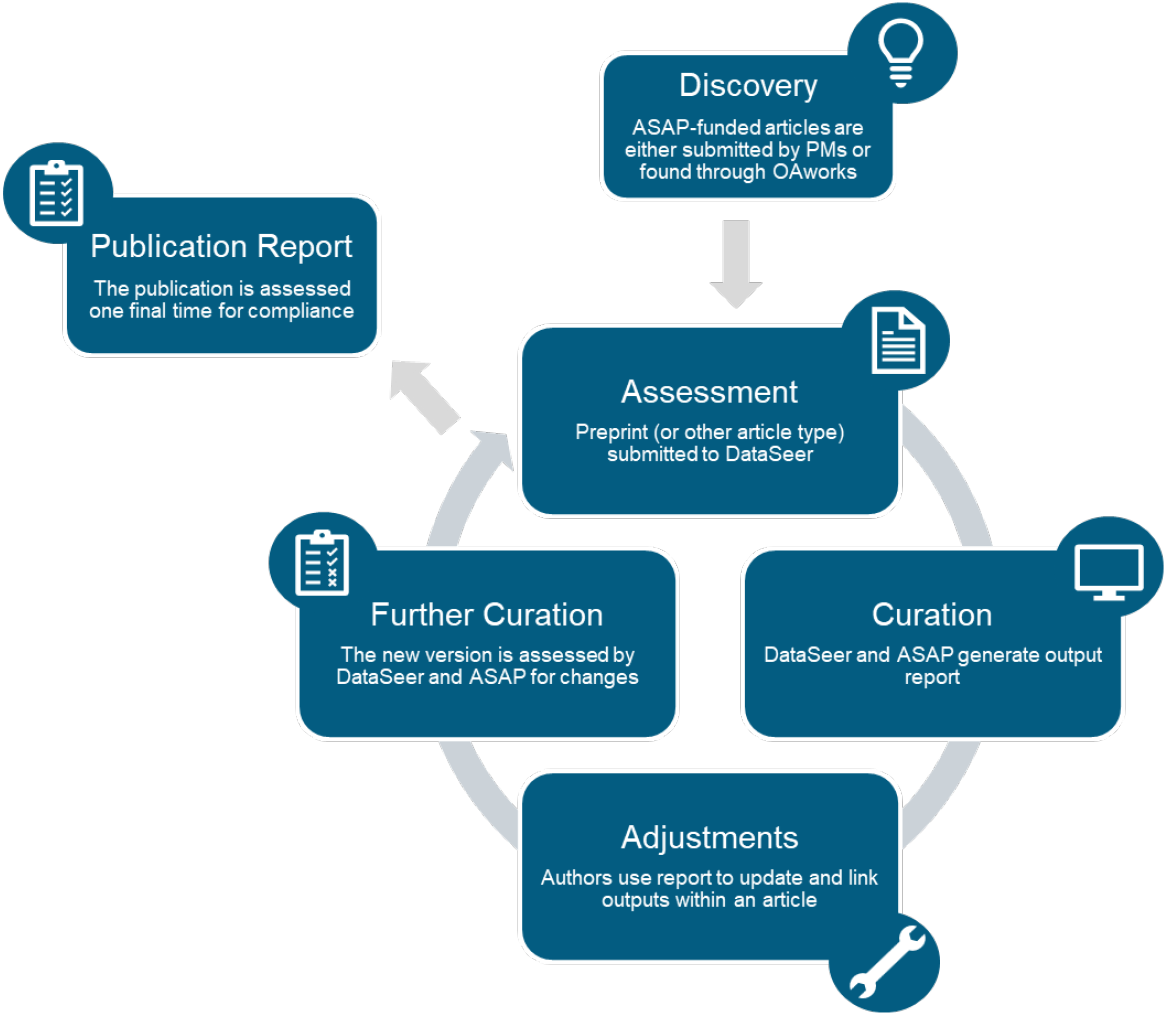
Workflow schematic the compliance workflow for ASAP grantees. ASAP-funded articles are discovered by two main mechanisms: either by Project Managers (PMs) or through OAworks, and then submitted to DataSeer on a rolling basis. Once received, DataSeer generates a compliance output report, which is checked by ASAP staff, and then shared with the authors of the article. Authors use the compliance report to understand what research outputs are not properly cited and recommendations for proper citation. After the article is revised, it is resubmitted to DataSeer and curated again. The process of assessment - curation - adjustment can be repeated until all research outputs are properly cited. Finally, when the article is ready for publication, the article is assessed one final time for compliance.

### Standardized Research Output Compliance Rules

ASAP and DataSeer have been developing consistent, enforceable rules for research outputs based on available [FAIR standards](https://www.go-fair.org/fair-principles/). FAIR rules are well-established for data and generally for code but are not yet as concrete for other trackable research outputs available in academic publications. For example, the rules for citing the reuse of software packages remain subjective despite the existence of research resource identifiers, RRIDs. Often creators of software ask users to cite a specific manuscript rather than an RRID.

Due to the complexity of the landscape, we count any object as properly shared if an object has specific, functioning, stable identifier(s). Table 1 lists the definitions of each of ASAP’s tracked research outputs in a manuscript and Table 2 displays the criteria for appropriately shared research outputs as per ASAP guidelines. However, there are several exceptions within each research output type that continue to be addressed by the community at-large and described briefly in the assessing impact section below. Importantly, this workflow allows for any future changes to be implemented at scale.

**Table 1.** Tracked Research Output Types and Identifier Acronyms

| Table 1. Tracked Research Output Types and Identifier Acronyms |  |
| --- | --- |
| Research Output Definitions |  |
| **Data Collection** | Data collection is an action where features of the real world (heights, reflectances, etc.) are captured by some device and become data points in a computer. A collection of data points about the same object or class of objects is a dataset. Reused data was generated in other publications or sources. |
| *Dataset Categories that currently are not required to be included in the final dataset deposition* |  |
| *Quality Control (QC) datasets* | Raw data from testing a sample of the output is not required, but measurements are recommended to be included to increase faith in results presented. All data types can appear in this category. |
| *Confirmatory datasets* | Intermediate steps that produce the final dataset can be treated exactly the same as QC datasets. Examples of Confirmatory datasets include many assays, chromatography, genotyping, confirmatory sequencing , etc. |
| *Representative Media datasets* | Visual and other datasets that sometimes require an enormous amount of space to store are not currently required to be included in repository uploads. Representative datasets can be new or re-used datasets. Examples of these data types include image, video, sound. |
| **Code** | Any usage of command-line software indicates that the authors must have custom scripts (which they wrote to operate the software), and these scripts need to be shared. Reuse of code is rare, but exists and requires adequate citation. |
| **Software** | Programs including software, packages, sites, databases, and other resources. New software is any program generated with ASAP funds. Appropriate citations are required for any use of existing software in the analysis of data or running simulations. |
| **Lab Materials** | Stable resources including organisms, cell lines, plasmids, and antibodies. |
| **Protocols** | Subsections within the “Methods” of an article excluding lists and statistical analyses. |
| **Acronyms** |  |
| **DOI** | Digital Object Identifier |
| **URL** | Uniform Resource Locator |
| **RRID** | Research Resource Identifier |
| **PID** | Persistent Identifier |

**Table 2.** Identifier Requirements for Appropriately Shared Research Outputs.

|  |  |  |  |  |
| --- | --- | --- | --- | --- |
|  | **Data** | **Code and Software** | **Lab Materials** | **Protocols** |
| **New** | URL or DOI | DOI (RRID if software) | RRID | URL or DOI |
| **Re-used** | URL or DOI, if URL include date queried | Version, URL, RRID (if available) | Source, Catalog Number, RRID (if available) | Citation, URL, DOI, or Kit Number |

## Assessing Impact: Compliance reports and measuring behavior change

Throughout ASAP’s short tenure, we aimed to continuously evaluate and adjust our infrastructure and methodology to better serve our network. As such, we sought to understand the feasibility, ease, impact, and improvement of our open science policies as they were put in practice with CRN teams.

### Inclusion Criteria

Compliance reports from DataSeer track and evaluate baseline compliance (prior to any intervention) and change over time as the research team understands their output sharing mistakes and amend these issues. To this end, we assessed output sharing in the first and second version of manuscripts coming through the DataSeer workflow (Figure 1). Because we continually refined the research output ruleset document through February 2022, we restricted evaluation to articles that were submitted to DataSeer after March 1, 2022 (termed first version). The subsequent submission (termed second version) were articles submitted to DataSeer at least two business days ***after*** the first version. This criterion was included as sometimes a draft manuscript was received the day it was also being uploaded to a preprint server, which appeared online 2 days later. Within this criterion, there were 19 different articles that had a first and second version assessment through our system between March 1, 2022, and October 1, 2022.

We then tallied the number of objects identified in the first and second version of the manuscripts based on ASAP’s standards (Table 2) for each of the newly generated and re-used research output types (datasets, code and software, lab materials, and protocols). We report both the proportions of output sharing by type alone with total numbers of objects identified by type as quantities of outputs can change between article versions. Please see Table 3 and Figure 2 for a summary of the overall results by object type.

**Table 3.** Summary statistics for assessing compliance over time.

| **Table 3. Summary statistics for assessing compliance over time.** |  |  |  |  |  |
| --- | --- | --- | --- | --- | --- |
|  |  | **Version 1** | **Version 2** | **Version 1** | **Version 2** |
|  | Number of papers referencing an output type | Total objects shared across all papers / Total objects identified across all paper | Total objects shared across all papers / Total objects identified shared across all papers | Average percentage shared per paper | Average percentage shared per paper |
| **New Datasets** | 16 | 13/108 (12%) | 95/137 (69%) | 29.25% | 62.05% |
| **New Code & Software** | 16 | 2/31 (6 %) | 21/32 (65%) | 12.50% | 43.75% |
| **New Lab Materials** | 5 | 0/424 (0%) | 28/428 (6%) | 0.00% | 1.81% |
| **New Protocols** | 13 | 12/89 (13%) | 52/99 (53%) | 16.67% | 59.56% |
| **Re-used Datasets** | 11 | 61/71 (86%) | 74/78 (95%) | 58.65% | 71.59% |
| **Re-used Code & Software** | 19 | 21/209 (10%) | 125/216 (58%) | 11.74% | 50.98% |
| **Re-used Lab Materials** | 13 | 48/281 (17%) | 156/387 (40%) | 13.02% | 40.33% |
| **Re-used Protocols** | 10 | 24/49 (49%) | 49/52 (94%) | 56.05% | 83.33% |
| *There were a total of 19 article pairs examined for this analysis. If a paper did not contain an object type in either the version 1 or version 2 assessment it was excluded from the analysis for that output type.* |  |  |  |  |  |

**Figure 2:**
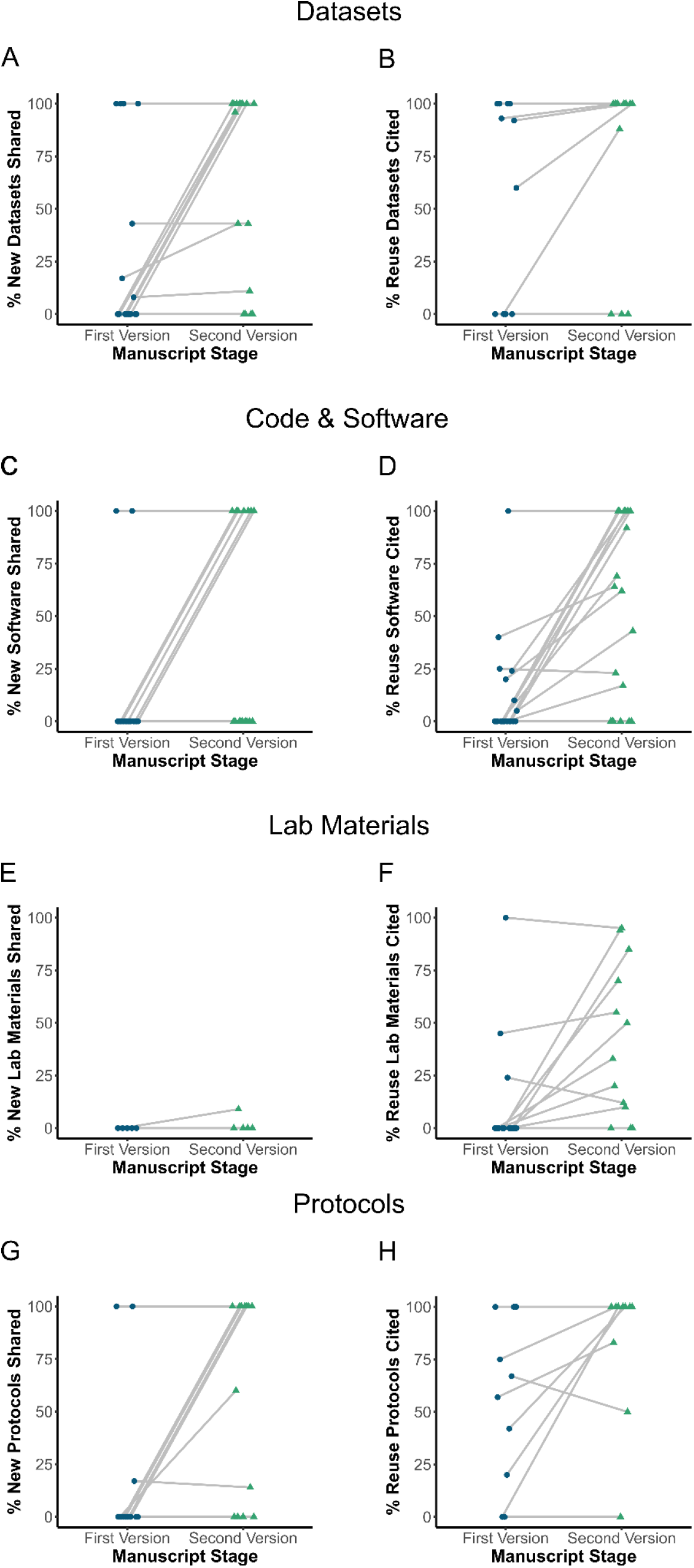
Compliance review increases the sharing of new and re-use research outputs. We assessed compliance with ASAP open science policy at two stages within the lifetime of ASAP-funded research articles: the first version submitted to ASAP for initial review, and the subsequent version following said review. Compliance was assessed by measuring the percentage of novel and re-use research outputs (datasets, code/software, lab materials, protocols) shared or cited in both versions. A total of sixteen manuscripts (16 research articles, 32 unique documents) were tracked in our analysis. Within each panel, the y-axis represents the percentage of shared research outputs, and the x-axis represents the two stages of manuscript development. Blue dots and green triangles represent unique manuscripts in the first and second versions. First and second versions of research articles are connected via gray lines **(a)** Percentage of new datasets shared across stages **; (b)** Percent reuse datasets cited across stages **(c)** Percentage of new code software shared across stages; **(d)** Percentage of reuse code and software cited **(e)** Percentage of new lab materials shared across stages; **(f)** Percentage of reuse lab materials cited across stages **(g)** Percentage of new protocols shared across stages; **(h)** Percentage of reuse protocols cited across stages

### Baseline Compliance

The baseline evaluation of whether a newly generated research output was appropriately linked in a first version manuscript was low on average ranging from 0% to 13% depending on output type (Figure 2, Table 3). There was a variability of compliance observed, with some authors at 100% compliance for certain object types in the initial version assessment. However, these instances were an exception rather than the rule (Figure 2).

For re-used outputs, citation of research outputs in the first manuscript version was higher compared to newly generated outputs, ranging on average from 10% to 86% of objects shared depending on output type. Authors were more likely to inaccurately cite lab materials (17% of objects identified were shared across all papers) and software packages (19% of objects identified were shared across all papers) compared to the other output types (Figure 2, Table 3). This could primarily be attributed to lack of consensus and education within the field on how to accurately cite software and lab materials with persistent identifiers in manuscripts. Most members within our network were not aware of Zenodo’s capabilities to sync up with GitHub to provide a digital object identifier (DOI) or what a Research Resource Identifier (RRID) was, let alone where to find associated RRIDs for resources. Based on this observation, we began to educate our project managers about DOIs and using the SciCrunch database to look up RRIDs.

### Compliance over time

After the first version of the article was submitted to DataSeer, teams received a compliance report that summarized action items needed for each output type to achieve compliance per ASAP policies. When evaluated in aggregate, all output types show an improvement in their sharing status in the second version of the manuscript (Figure 2, Table 3). This suggests that these reports were overall effective in assisting authors in appropriate linkage of newly generated research outputs to their manuscripts.

#### Dataset sharing and reuse

Most individual articles showed better sharing of newly generated datasets (percentage increasing on average from 12% to 69%) from first to second version (Figure 2A, Table 3). When asked why all datasets were not shared for a specific manuscript, the most common response was that the dataset in question had not been generated by the ASAP grantee and the grantee could not control the actions of collaborators who were not funded by ASAP. However, half of the articles (8 of the 16) assessed showed a complete shift to sharing all datasets in the second version, implying that once authors were made aware, they were willing to curate and deposit the identified datasets in appropriate repositories. Another source of confusion around datasets was about whether supplemental files could be considered an appropriate deposition for the dataset linked to the article. We decided to include datasets shared in supplemental files for this initial analysis, and are developing additional training material for future manuscripts on why deposition to a FAIR repository is preferred to this approach.

For re-used datasets, compliance on average was higher, jumping from 86% to 95% in the second version (Figure 2, Table 3). The use case for when re-used datasets were not accurately cited were due to circumstances outside an author’s control. In these instances, we asked the authors to provide as much additional information as possible, such as the contact of the individual who owned the data and the date the dataset was queried.

#### Code & Software sharing and reuse

For the purposes of this discussion, we will use the term software liberally to apply to both code and software outputs. When assessing software in the second version of manuscripts, there was a strong upward trend toward more software shared, this trend is seen in both new and reused software outputs (Figure 2C, D). In the case of new software shared, many articles shifted from no software shared to all software shared (Figure 2C). Software that was reused did show a general upward trend, but the individual articles are more variable in their compliance (Figure 2D). Often, this has to do with the lack of clear guidelines on the best way to cite existing software. For example, software developers may ask that the citation be made in the form of a specific publication versus an RRID or a URL associated with the instance in question. Moreover, not all existing software has been registered with a permanent identifier in the SciCrunch database or other repositories such as Zenodo. There is a hesitancy to register software on someone else’s behalf and doing so may also create multiple RRIDs for the same software instance. Authors often do not understand the value of permanent identifiers over publication citations and version numbers.

#### Lab resource sharing and reuse

The greatest challenge for authors was with registering new lab materials generated from the manuscript. Even in the second version of manuscripts, only 6% of newly generated materials had an RRID associated with the output (Figure 2E). This is largely due to three main issues. First, it takes time to get a resource deposited and available for distribution in a registry and many authors did not realize that one can pre-register a resource. Second, there is general confusion as to how an RRID should be registered, as different resource types are governed by different agencies that have different procedures (antibodies are handled separately from cell lines, for example). Third, there are certain stable resources that currently do not have registering bodies that can mint RRIDs, such as newly generated gene probes or compounds and there is no clear framework outlined for using patent numbers or other isolated identifiers for these use cases.

#### Protocol sharing and reuse

There was a strong upward trend toward sharing protocols (Figure 2G, H, Table 3). The percent shared jumped from 13% in the first version to 53% in the second version on average. During our outreach, we learned that the biggest barrier to sharing methods was that authors were worried about plagiarism and didn’t understand the rationale for why we were requiring the methods sections to link out to a recipe style registered protocol. Most believed that the description in the methods section of a manuscript was enough information. To help teams, ASAP provided information on how protocols are not copyrighted material, emphasizing that credit should still be given but anyone can upload a protocol (if it was not a trademark secret) regardless of who generated it. Additionally, ASAP staff shared lessons learned from the Cancer Reproducibility Project which proved to be a great motivator in bringing individuals on board with registering with platforms like protocols.io and sharing protocols (Errington et al., 2021).

#### Other Considerations

As we continue to evaluate our ability to link research outputs, we are also turning towards assessing the completeness of the metadata around the data deposition. To help train our network, ASAP developed a [checklist for repository deposition](https://zenodo.org/record/7405544#.Y_9Unj1BxPY), explaining the components to consider and the rationale for why it mattered.

Software and other key resources used in the generation of figures, tables, and statistics are listed in Table 4 in the Data and Code Availability section

**Table 4.** Key Resources

| Table 4. Key Resources |  |  |  |
| --- | --- | --- | --- |
| Version | Software Name | URL | RRID |
| v0.0.1 | DataSeer | https://dataseer.ai/ | RRID:SCR_023027 |
| 1.0.10 | dplyr | https://dplyr.tidyverse.org | RRID:SCR_016708 |
| 3.4.0 | ggplot2 | https://ggplot2.tidyverse.org | RRID:SCR_014601 |
| n/a | Google Sheets | https://www.google.com/sheets/about/ | RRID:SCR_017679 |
| n/a | OA.Report | https://oa.report/ | RRID:SCR_023288 |
| n/a | Protocols.io | http://protocols.io/ | RRID:SCR_010490 |
| 4.1.1 | R Project for Statistical Computing | https://www.r-project.org/ | RRID:SCR_001905 |
| n/a | SciCrunch | https://scicrunch.org/ | RRID:SCR_003115 |
| n/a | ZENODO | https://zenodo.org/ | RRID:SCR_004129 |

## Roadmap for the future

We have observed two main barriers to compliance. First, there is a lack of a community framework on how research outputs should be registered, deposited, and cited. Second, there is a lack of education on the best open science practices as they stand today. While it is expected that open science standards may change in the coming years as the landscape is evolving, it will be important to note the current best practices for a particular point in time to ensure a consistent message and framework upon which compliance monitoring can be built. ASAP aims to contribute to the open science community by educating our growing network (currently over 1000+ individuals). To aid with training, our project managers serve as open science ambassadors for their team. The project managers have monthly training to stay current with best practices in open science and ASAP requirements as well as share roadblocks with the ASAP open science team.

As more funding bodies and institutions embrace open research, there are key actions that, if taken collectively, would ignite rapid culture change and assist with compliance with the emerging landscape of open research goals and policies:

1. **Align policies and offer direct incentives** for collaboration, transparency, and reproducibility in research communication.
2. **Define compliance and establish open standards** for tracking and measuring open research practices so that it is clear when compliance is reached.
3. **Establish common best practices and standards** including repository use, appropriate persistent identifiers (PIDs) to use depending on research output type, and clear instructions on how to efficiently share and log outputs.
4. **Invest in infrastructure** that helps existing repositories become FAIR compliant, creates pathways for appropriate PID assignment, removes PID delays, and standardizes compliance metrics.
5. **Normalize the schema for outputs**, an extensible publicly owned research output management schema used across all infrastructures to prevent the fractured metadata landscape that plagues the published record today.
6. **Automate and streamline sharing** by depositing article supplementary materials into FAIR repositories with PIDs assigned, detecting datasets that haven’t been shared, linking deposits to ORCIDs and articles, updating outputs based on connected publications.
7. **Pool these actions** across funding bodies, institutions, and publishers so that they can scale.

Our analysis shows that most authors are willing to comply with open science practices if the policies and requirements are clearly outlined and support is provided through the project manager role. ASAP requires each team to hire a project manager with PhD credentials, whose role is to educate the team members and assist with compliance support around open science policies. Many of the current barriers to persistent identifiers are due to lack of awareness surrounding best practices and the fragmented infrastructure for registering research outputs in multiple locations. By working with other funders as well as institutions and research communities, ASAP hopes to help influence a widespread uptake of collaborative and open research practices and to contribute to a shared knowledge base on how best to establish this as the norm for the coming years.

## Data and Code Availability Statement

All data including submission output proportion data and the Compliance Rules used are available along with the analysis code on Zenodo, https://doi.org/10.5281/zenodo.7504034

The ASAP Repository Checklist helps to screen outputs for proper sharing and metadata completeness https://doi.org/10.5281/zenodo.7405544

The ASAP Blueprint for Collaborative Open Science v1 a comprehensive report on how ASAP has worked towards its goals to date and includes all associated materials, templates, and schemas 10.5281/zenodo.6979998

## Acknowledgements

The authors thank the Sergey Brin Family Foundation for funding and supporting the Aligning Science Across Parkinson’s initiative and The Michael J. Fox Foundation for their indispensable expertise in implementing the ASAP initiative. A special thanks to Lindsey Riley for her feedback and review of the draft manuscript. The authors also thank Sarah Greene of Rapid Science who helped formulate ASAP’s initial open science policies, Alyssa Yong from DataSeer.AI, Antia Bandrowski from SciCrunch and Joe McArthur from OA.works for helping us shape our open science ecosystem.

This research was funded by Aligning Science Across Parkinson’s through the Michael J. Fox Foundation for Parkinson’s Research (MJFF). For the purpose of open access, the author has applied a CC-BY public copyright license to the Author Accepted Manuscript version arising from this submission.

---

**Table 1. Definitions for each research output type and acronym definitions for different identifiers**

**Table 2. Requirements for appropriately shared research outputs**

**Table 3. Summary statistics for assessing compliance over time**

**Table 4. Key Resources and RRIDs**

## Notes

https://doi.org/10.5281/zenodo.7504034

https://doi.org/10.5281/zenodo.7405544

### Competing Interest Statement

The authors have declared no competing interest.

### Summary of Updates

This version of the manuscript was revised to update the author list order

